# Integrative Proximal-Ubiquitomics Profiling for Deubiquitinase and E3 Ligase Substrate Discovery Applied to USP30

**DOI:** 10.1101/2024.10.07.616967

**Authors:** Andreas Damianou, Hannah B.L. Jones, Athina Grigoriou, Iolanda Vendrell, Simon Davis, Benedikt M. Kessler

**Affiliations:** Target Discovery Institute, Centre for Medicines Discovery, Nuffield Department of Medicine, University of Oxford, Roosevelt Drive, Oxford OX3 7FZ, UK; Chinese Academy for Medical Sciences Oxford Institute, Nuffield Department of Medicine, University of Oxford, Roosevelt Drive, Oxford OX3 7FZ, UK

**Keywords:** APEX2, proximity labelling, deubiquitinase, ubiquitin specific protease 30, USP30, LETM1, mass spectrometry, proteomics

## Abstract

Increasing interest in deubiquitinases (DUBs) and ubiquitin E3 ligases as drug targets to modulate critical molecular pathways in disease is driven by the discovery of specific cellular roles of these enzymes. Key to this is the identification of DUB or E3 ligase substrates. While global cellular ubiquitination changes upon perturbation of DUB/E3 ligase activity can be studied using mass spectrometry-based proteomic methods, these datasets include indirect and downstream ubiquitination events. To enrich for direct substrates of DUB/E3 ligase enzymes, we have combined proximity-labelling methodology (APEX2) and subsequent ubiquitination enrichment (based on the K-ε-GG motif) to form a proximal-ubiquitome workflow. We have applied this technology to identify altered ubiquitination events in the proximity of the DUB ubiquitin specific protease 30 (USP30) upon its inhibition. We show ubiquitination events previously linked to USP30 on TOMM20 and FKBP8 and the previously undescribed candidate substrate LETM1, which is deubiquitinated in a USP30-dependent manner.

## Introduction

Ubiquitin (Ub) constitutes a highly conserved protein found across all eukaryotic organisms.^1^ Ub can be activated and covalently attached to a target protein via the successive action of E1, E2, and E3 enzymes, forming a cascade that culminates in a post-translational modification. This intricate process plays an instrumental role in a myriad of cellular functions, encompassing protein homeostasis, signal transduction, DNA replication, and transcriptional regulation.^2^

E3 ligases and deubiquitinating proteases (DUBs) are specialist enzymes that add and remove covalent ubiquitin modifications to and from target substrates. Both enzyme families are essential in the maintenance of Ub homeostasis. This dynamic balance is integral to normal cellular operation, and can alter the fate of Ub-modified proteins.^3^ Imbalances in the ubiquitination of proteins have been implicated in various pathological conditions, including cancer and neurodegenerative diseases.^4,5^ Therefore, the identification of E3 ligase/DUB substrates not only enriches our understanding of their biological roles but also opens avenues for the exploration of these enzymes as potential therapeutic targets.^6^

Several methodologies have been previously employed to identify potential substrates of E3 ligases that could also be translated to identify DUB substrates. One category involves fusion techniques, such as UBAITS,^7^ TULIP,^8^ and TULIP2,^9^ which use overexpressed Ub-E3 ligase fusions to trap and purify ligase-substrate adducts, often under denaturing conditions. Another category includes *in vitro* techniques such as E2-dID, orthogonal Ub transfer (OUT),^10,11^ and microarray assays, which employ engineered enzymes or biotin-labeled Ub conjugates to target specific substrates. Finally, UbPOD^12^ and BioE3^13^ are approaches that use tagged ubiquitin as a biotinylation substrate for an E3 ligase conjugated to a proximity labelling enzyme. While no single method is universally applicable, each has proven effective for discovering substrates for specific E3 ligases.

While these methods enrich for interactors of E3 ligases, they do not provide information on the ubiquitination sites on these potential substrates. The identification of specific ubiquitination/deubiquitination sites is critical for interpreting the biological meaning of the ubiquitination events altered by an E3 ligase/DUB, and for utilising the ubiquitination site as a biomarker in the case that an E3 ligase/DUB is therapeutically targeted. Additionally, altered ubiquitination states on proteins that are E3 ligase/DUB interactors increases the confidence that these hits may be direct substrates.

Among the most established approaches to identify ubiquitin sites is the enrichment of the Ubiquitin Remnant Motif (K-ε-GG) left behind at the sites of lysine ubiquitination after protein trypsinization, followed by immunoprecipitation and subsequent mass spectrometry analysis.^14,15^ The application of targeted knockouts, catalytically inactive mutants or of specific small molecule inhibitors, in conjunction with the K-ε-GG technique, can accelerate the identification process of DUB or E3 ligase substrates from specific alterations observed in the cellular ubiquitome.

However, the K-ε-GG method comes with limitations. One notable drawback is that many altered ubiquitination events may result from downstream secondary effects upon the reduced activity of a DUB or E3 ligase. Whilst these events are of interest, they are not all reflective of direct enzyme substrates and activity.

In the last decade, innovative proximity labelling approaches, such as BioID^16^ and APEX^17^, have been developed to enhance the identification of proteins within the micro-environment of a protein of interest. Proximity labelling has previously been successfully integrated with phospho-enrichment strategies to investigate spatially referenced proximity phosphoproteomics.^18–20^ In this study, we show that the effective combination of proximity labelling followed by post-translational modification enrichment can also be applied to the study of ubiquitination. This allows for the capture of protein ubiquitination states that are specifically localised to DUBs/E3 ligases. Application of the technique has the potential to enhance the resolution and accuracy of substrate identification of a DUB or E3 ligase upon the perturbation of its activity.

Ascorbate peroxidase-2 (APEX2) was considered favourable as a proximity labelling enzyme for this methodology due to the speed of the biotinylation reaction occurring on the second timescale. This labelling snapshot enables the capture of an acute cellular protein ubiquitylation profile upon DUB inhibition/genetic depletion, without time-dependent convolution from the dynamic ubiquitination process. Additionally, APEX2 favours the biotinylation of tyrosine residues, meaning that it does not interfere with ubiquitination events occurring on lysine residues. Other proximity labelling approaches, such as BioID and TurboID, should not be applied to this methodology due to the long timescale of the labelling reaction and the biotinylation of lysine residues.

USP30 was selected as a model DUB for the development of this methodology due to its potential as a drug target, with the knockout of USP30 resulting in increased turnover of mitochondria via mitophagy.^21^ The process of mitophagy could be targeted for multiple therapeutic applications, such as for the treatment of Parkinson’s disease.^21,22^ Therefore, further insight into the role USP30 plays in the process of mitophagy, and potential biomarkers resulting from its inhibition are highly relevant. Additionally, previous characterisation of the outer mitochondrial membrane (OMM) localisation of USP30, and ubiquitomics studies of the activity of USP30 enable cross-validation of the proximal-ubiquitome of USP30 to existing datasets.^22–24^

Here, we present the development of a novel workflow, combining highly sensitivity APEX2 proximity labelling with a subsequent K-ε-GG immunoprecipitation to elucidate the proximal-ubiquitome of a target DUB (USP30), both in the presence and absence of an USP30 inhibitor, compound ‘**39**’, a previously described potent, cell permeable, and selective USP30 inhibitor.^25–27^ The proximal-ubiquitomics methodology gives a unique insight into the role of the USP30 in process of mitophagy. Potential direct USP30 substrates are uncovered through the application of the methodology, and subsequent validation provides evidence for USP30 enacting on the ubiquitination state of proteins considered critical to mitochondrial function and turnover. We focus on validating a novel previously undescribed potential USP30 substrate – leucine-containing zipper and EF-hand transmembrane protein 1 (LETM1), an essential protein described to be involved in various critical functions, including mitochondrial ion transport, regulation of mitochondrial volume, energy metabolism, and maintenance of mitochondrial morphology.^28–31^

## Results

### Proximal-ubiquitomic methodology optimization

Each aspect of the workflow applied in Figure 1 was optimised to ensure efficient capture of ubiquitylated proteins in the proximity of USP30 (Figure 2a). This is critical for the adaptation of the methodology to different systems. In this model USP30-APEX2 system, the conjugated enzymes were stably expressed in HEK293 cells with a low expression promoter (PGK).^32^ We observed a moderate 3.8-fold increase in USP30 expression as compared to that of its endogenous expression levels (Figure 2b & S1a). Maintained USP30 activity was confirmed using a DUB activity-based probe (ABP), consisting of ubiquitin with a propargylamine warhead and a haemagglutinin tag (HA-Ub-PA). The ∼10 kDa molecular weight increase of USP30-APEX2 in the presence of HA-Ub-PA indicates that the active site of USP30 is still able to bind ubiquitin and the active site cysteine is still reactive (Figure 2c & S1a). This ABP is broadly reactive with the majority of cysteine-active DUBs, and so can be used to check retained activity if the proximal-ubiquitomics methodology is applied to other DUBs.^33^ The maintained localization of USP30-APEX2 to the OMM was confirmed via TOMM20 co-localisation microscopy (Figure 2d) and digitonin membrane isolation (Figure S1b).

**Figure 1.**
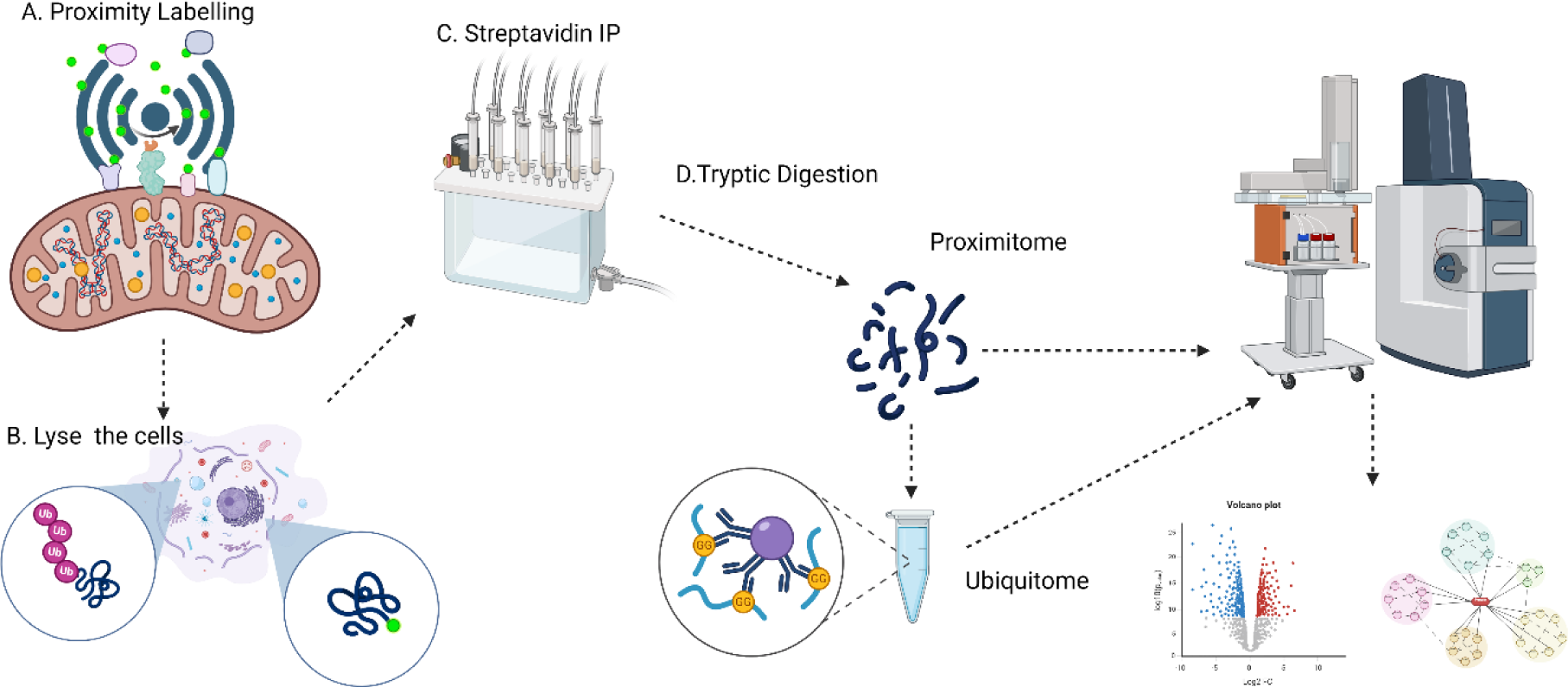
Integrated proximal-ubiquitomics methodology workflow. Proximity labelling biotinylates the microenvironment of a DUB/E3 ligase of interest. The biotinylated proteins are then pulled down with streptavidin on the protein level to encompass ubiquitination events proximal to the biotinylation site. Biotinylated proteins are then digested with trypsin and ubiquitination events are enriched using a K-ε-GG antibody. Ubiquitylated proteins from the proximity of the enzyme of interest are then quantified by LC-MS/MS. Figure created using www.biorender.com.

**Figure 2.**
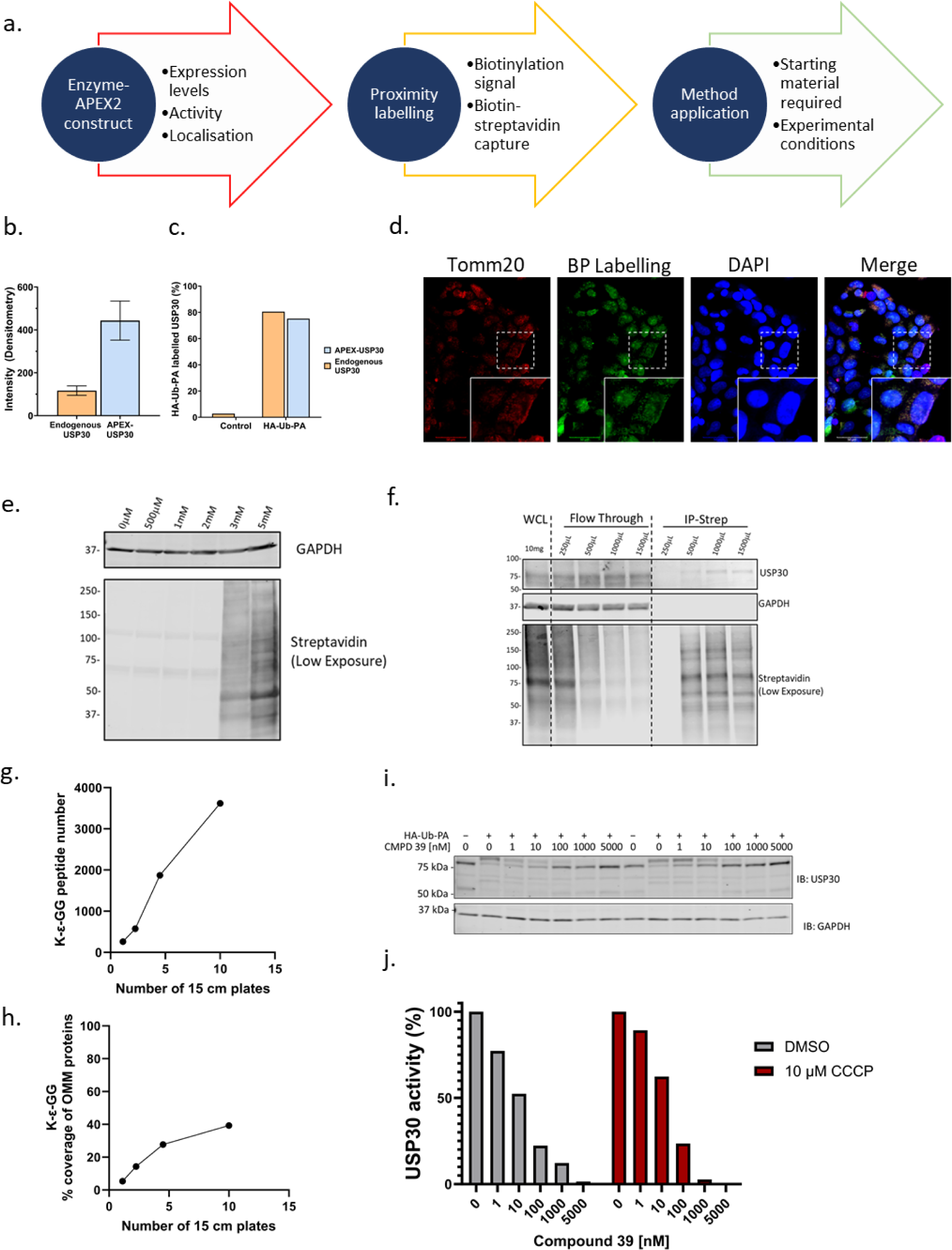
Optimization workflow for the proximal-ubiquitomics methodology. **a.** Scheme of experimental optimization strategy. **b.** Densitometric intensity of HEK293 endogenous USP30 and APEX2-USP30 (N=3 quantification of separate bands from 1 sample). **c.** Percentage of endogenous and APEX2 conjugated USP30 labelled by HA-Ub-PA. **d.** Immunofluorescence of TOMM20, biotin-phenol (BP) and DAPI staining after USP30-APEX2 biotinylation in HEK293 cells. **e.** Biotin-phenol concentration dependence of USP30-APEX2 biotinylation in HEK293 cells. **f.** Streptavidin pulldown optimization after USP30-APEX2 biotinylation with 5 mM biotin-phenol. **g.** Number of GlyGly peptides identified from the proximal-ubiquitomics methodology with different amounts of input material. **h.** Percentage of outer mitochondrial membrane (OMM) Mitocarta 3.0 proteins identified from the proximal-ubiquitomics methodology with different amounts of input material. **i.** HA-Ub-PA labelled USP30-APEX after a 6hr incubation of the USP30-APEX2 HEK293 cells with the indicated concentration of compound 39, with and without 10 µM CCCP. **j.** Normalised densiometric quantification of USP30-APEX2 HA-Ub-PA labelling and inhibition after a 6hr incubation of the USP30-APEX2 HEK293 cells with the indicated concentration of compound 39, with and without 10 µM CCCP.

After confirming that the properties of the DUB-APEX2 conjugate are reflective of the endogenous DUB, the APEX2 biotinylation labelling efficiency was optimised, to maximise the capture of proximal proteins. Incubating cells with a concentration range of biotin demonstrated that a minimum of 5 mM biotin was necessary for efficient protein biotinylation (Figure 2e). In previous studies, biotin incubation with cells was performed for 30 minutes at 37 °C with increased incubation not resulting in increased labelling, indicating the incubation time is sufficient for full dispersal of the biotin throughout the cells.^34^ With the maximal biotinylation conditions, the volume of streptavidin beads then needs to be optimised to achieve efficient capture of all biotinylated proteins. By assessing the input, flow-through and elution of the biotin streptavidin pull down, we maximised the capture capacity of the streptavidin beads for this system with 1 mL of beads / 10 mg of input protein (Figure 2f).

Once proteins proximal to USP30 were biotinylated by APEX2 and pulled down using streptavidin, the ubiquitination sites of these proteins were enriched for LC-MS/MS identification. This was achieved by trypsinizing the proteins and immunoprecipitating the K-ε-GG remnant motif that remains after the trypsinization of proteins reflecting ubiquitination, Neddylation or ISGylation sites. However, under most normal physiological conditions, the source of the majority of K-ε-GG remnant sites is ubiquitination (>94 %).^35^ Without K-ε-GG pulldown, we did not detect ubiquitination sites in the USP30 proximity labelling (data not shown), justifying the necessity for a subsequent enrichment step. The recommended amount of protein material for K-ε-GG enrichment is 1-2 mg.

Measurement of protein/peptide levels is challenging in the workflow due to interference from the presence of anti-oxidants in the lysis buffer. Additionally, proteins such as streptavidin and trypsin obscure the absolute protein quantitation after biotinylation elution and the peptide quantitation prior to the K-ε-GG enrichment, respectively. Therefore, material for this experimentation was optimised for the system based on the number of 15 cm plates at confluence used as starting material. When performing K-ε-GG enrichments without proximity enrichment the number of identified K-ε-GG sites identified can vary greatly, depending on the source material, experimental conditions and mass-spectrometry methodology applied.^36,37^

In the case of the proximal-ubiquitomics workflow presented here, we saw a linear increase in the number of K-ε-GG sites with increasing input material, up to 4000 sites with 10 x 15 cm plates (Figure 2g). However, the coverage of OMM localised proteins that were identified as ubiquitinated begins to plateau between 5 - 10 x 15 cm dishes of input material (Figure 2h). Thus, 5 x 15 cm dishes per condition were selected as the optimal input material for this system to capture ubiquitination events proximal to USP30. The abundance of USP30 has been reported to be in the bottom 10 % of ∼8000 proteins identified in a HEK293 proteome (PaxDB 5.0).^38,39^ With the use of a low expression promotor to mimic endogenous USP30 levels in this study, large amounts of input material were required for this model system. This may vary depending on expression levels of the protein of interest, and subsequently the amount of material recovered post streptavidin IP.

After optimization of the USP30-APEX2 system and proximal-ubiquitomics workflow, experimental conditions to elucidate potential USP30 substrates using compound **39** were optimized. To maximize ubiquitination events in the proximity of USP30, cells were treated with CCCP to depolarize the mitochondria and subsequently activate the PINK1/Parkin S65 phospho-ubiquitination pathway. Mitochondrial depolarization conditions have previously been shown to reveal increased OMM protein ubiquitination events as a consequence of USP30 knockout.^24^ In this HEK293 USP30-APEX2 system, maximal S65 phospho-ubiquitin was identified with 10 µM of CCCP cell treatment after 6 hours of incubation (Figure S1c).

The concentration of the small molecule compound **39** required for complete USP30 inhibition in the USP30-APEX2 overexpression system was assessed using the HA-Ub-PA ABP. After 6 hours of cellular incubation, full inhibition of HA-Ub-PA binding to USP30-APEX2 was identified with 5 µM of compound **39**, regardless of the presence of 10 µM CCCP (Figure 2i-j). From this optimization, 10 µM CCCP treatment +/− 5 µM of compound **39** was applied to identify acute ubiquitination events proximal to USP30 occurring as a consequence of USP30 inhibition on mitochondria undergoing the process of mitophagy.

### Optimised protocol reveals USP30 proximal ubiquitome enriched for OMM

The two conditions (CCCP treatment +/− 5 µM of compound **39**) were repeated in quadruplicate, with 5 x T175 flasks pooled per replicate. APEX2 biotin labelling was catalysed by the addition of H_2_O_2_ and quenched with multiple antioxidant washes. After lysis 5 % were taken for the ‘labelled proteome’, with the remaining material used as input for the biotin-streptavidin pulldown. Equal input and eluate levels for each streptavidin pulldown were checked by western blot (Figure S1d). From this pull down, 10 % of eluates were taken for the ‘proximitome’ and the remaining material was trypsinized and used as input for the K-ε-GG immunoprecipitation, forming the ‘proximal-ubiquitome’ (Figure 3a). Each step represents important information about the effect of USP30 inhibition as well as providing normalisation data to ensure ubiquitination events are not altered as a consequence of perturbed protein abundance or protein proximity to USP30 with compound **39** treatment.

**Figure 3.**
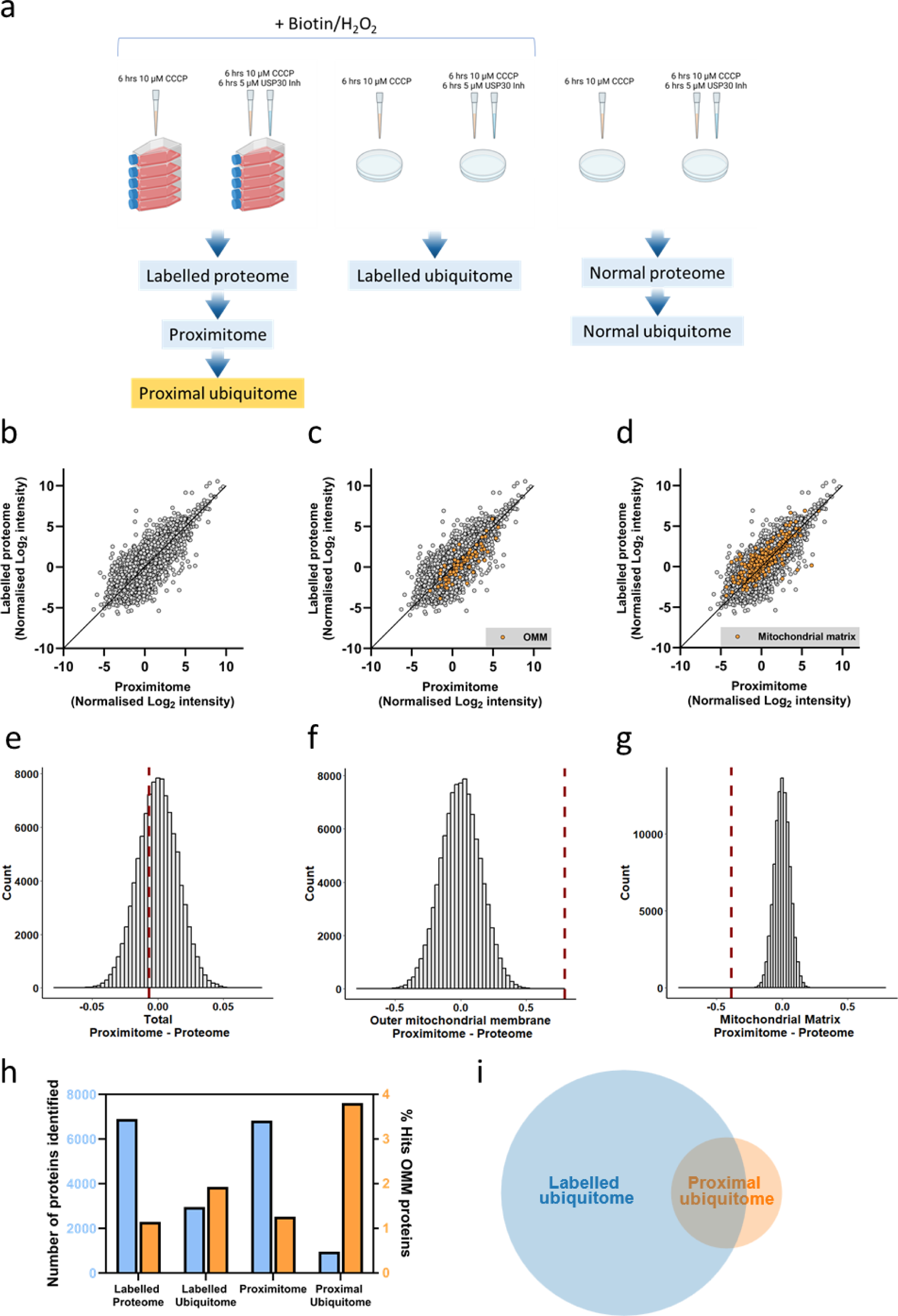
USP30 proximal ubiquitome enriched for outer mitochondrial membrane (OMM) proteins. **a.** Experimental design to compare the proximal ubiquitome of USP30 +/− compound **39,** with its corresponding proximitome, proteome and ubiquitome. A control proteome and ubiquitome without APEX2 biotin labelling were also included for comparison. **b-d.** The averaged Log_2_ intensity of proteins quantified in the proximitome *vs* proteome (with APEX2 biotinylation and CCCP treatment) with Mitocarta 3.0 annotated OMM proteins or MM proteins in orange. **e-g.** Permutation analysis – the dotted red line denotes the average of the X-Y intensity differences between the proximitome and the proteome of all proteins, outer mitochondrial membrane (OMM) proteins and mitochondrial membrane (MM) proteins. The distribution shows the randomized averages of these values over 100,000 permutations. **h.** The number of proteins identified in the biotin labelled proteome, labelled ubiquitome, proximitome and proximal ubiquitome vs. the % of Mitocarta 3.0 OMM annotated proteins. **i.** The overlap in K-ε-GG peptides identified in the labelled ubiquitome and proximal ubiquitome.

To compare the method of the proximal-ubiquitome to the classical ubiquitomics workflow we included a ‘labelled ubiquitome’ (N=3) with APEX2 biotinylation protein labelling, but without biotin enrichment by streptavidin. This allowed for the comparison of ubiquitination events occurring in the whole cell with those occurring in the proximity of USP30 in the same system. We also included a control proteome and ubiquitome (N=4) without APEX2 biotinylation labelling (‘normal proteome’ and ‘normal ubiquitome’), to assess the effect of the biotin/H_2_O_2_ treatment and subsequent APEX2 biotinylation reaction on protein and ubiquitination site identification/quantification (Figure 3a). The Log_2_ intensities for all datasets averaged coefficients of variations (CVs) below 10 %. The raw intensities showed higher CVs for the ubiquitomes compared to the proximitome/proteomes, with CVs for the proximal-ubiquitome comparable to the control ubiquitomes (Figure S2 a-f). Principal component analysis showed clustering of replicates based on presence or absence of USP30 inhibition (Figure S2 g-i).

To examine the effect of APEX2 biotin labelling on the proteome and ubiquitome of USP30-APEX2 expressing HEK293 cells, the labelled proteome was compared to the normal proteome, and the labelled ubiquitome was compared to the normal ubiquitome. There were significant changes to the intensity of proteins identified in the proteomes (Figure S3a), with String analysis^40^ identifying that the majority of proteins with significantly increased intensities upon APEX2 labelling are structural constituents of chromatin (Figure S3b). This may be attributable protein upregulation in response to DNA damage caused by the H_2_O_2_ treatment^41^ during the APEX2 labelling. No Mitocarta 3.0 annotated proteins were altered, indicating that the labelling of proteins with biotin does not affect the detection and intensity of the proteins anticipated to be proximal to USP30. There were also significant changes to the intensity of K-ε-GG peptides identified in the labelled ubiquitome when compared to the normal ubiquitome, including peptides originating from Mitocarta 3.0 annotated proteins (Figure S3c). The altered ubiquitome as a consequence of APEX2 catalysed H_2_O_2_ biotin labelling is a limitation of proximity labelling and highlights the importance of subsequent validation of hits identified with proximal-ubiquitomics in systems lacking APEX2 biotinylation process.

Proximity-labelling of USP30-APEX2 has a high background due to its exposure to a vast number of cytoplasmic proteins and the large number of proteins that bind as background during the streptavidin pull-down. However, proteins that are most frequently proximal to USP30 will include genuine USP30 interactors, and therefore potential substrates of USP30. OMM proteins are a key example of this, with the fixed positioning of these proteins likely to increase their USP30-APEX2 biotinylation frequency, and consequently their LC-MS/MS intensities. Additionally, previously identified USP30 dependent ubiquitylation events are localised to the OMM, such as TOMM20.^22,24,42,43^ To ensure the successful enrichment of proteins proximal to USP30-APEX2, the intensity of proteins annotated in Mitocarta3.0 as OMM *vs* mitochondrial matrix (MM) proteins were compared in the labelled proteome vs the proximitome (Figure 3b-d).

Permutation calculations were used to randomize the difference in X-Y scatter points in Figure 3b with the distribution of the average differences of the permutations plotted in Figure 3e. Here, the average difference in X-Y coordinates denoted by the red dashed line is within the normal distribution of the permutations, showing that the identified proteins are comparably intense between the proximitome and labelled proteome overall. When the same calculations are performed for the Mitocarta 3.0 annotated OMM proteins (Figure 3c-f) and MM proteins (Figure 3d-g), the average differences denoted by the red dashed line are significantly distinct from the randomized normal distribution, with the average difference never occurring in 100,000 permutations. These calculations show an enrichment of OMM proteins, and a reduction of MM proteins in the proximitome when compared to the labelled proteome. This confirms that the proximitome represents an enrichment of proteins that are situated within the proximity of USP30.

The enrichment of OMM proteins in the proximitome of USP30-APEX2 also translates through to the proximal-ubiquitome of the protein. The percentage of proteins linked to the OMM based on Mitocarta3.0 annotation is doubled in the proximal-ubiquitome compared to the control labelled ubiquitome (Figure 3h). Despite a reduced number of K-ε-GG modified peptides identified in the proximal-ubiquitome compared to the labelled ubiquitome, over 30 % of proximal-ubiquitome peptides were not identified in the labelled ubiquitome (Figure 3i). This indicates that proximal-ubiquitomics in this system enriches for a subset of ubiquitination events from the proximity of USP30, thus narrowing down potential substrates of the DUB.

### Application of proximal-ubiquitomics to identify novel substrates of USP30

Following the experimental outline in Figure 3a, the labelled proteome and the proximitome were assessed for protein intensity alterations occurring as a consequence of USP30 inhibition (Figure 4a-b). Although some proteins were significant in their alterations (denoted in orange), only 1 protein in the labelled proteome and no proteins in the proximitome were increased or decreased more than a Log_2_ fold change > 1 with USP30 inhibition. Minimal changes to the proteome as a consequence of USP30 inhibition or knockout have also been previously reported in the literature.^22,42^

**Figure 4.**
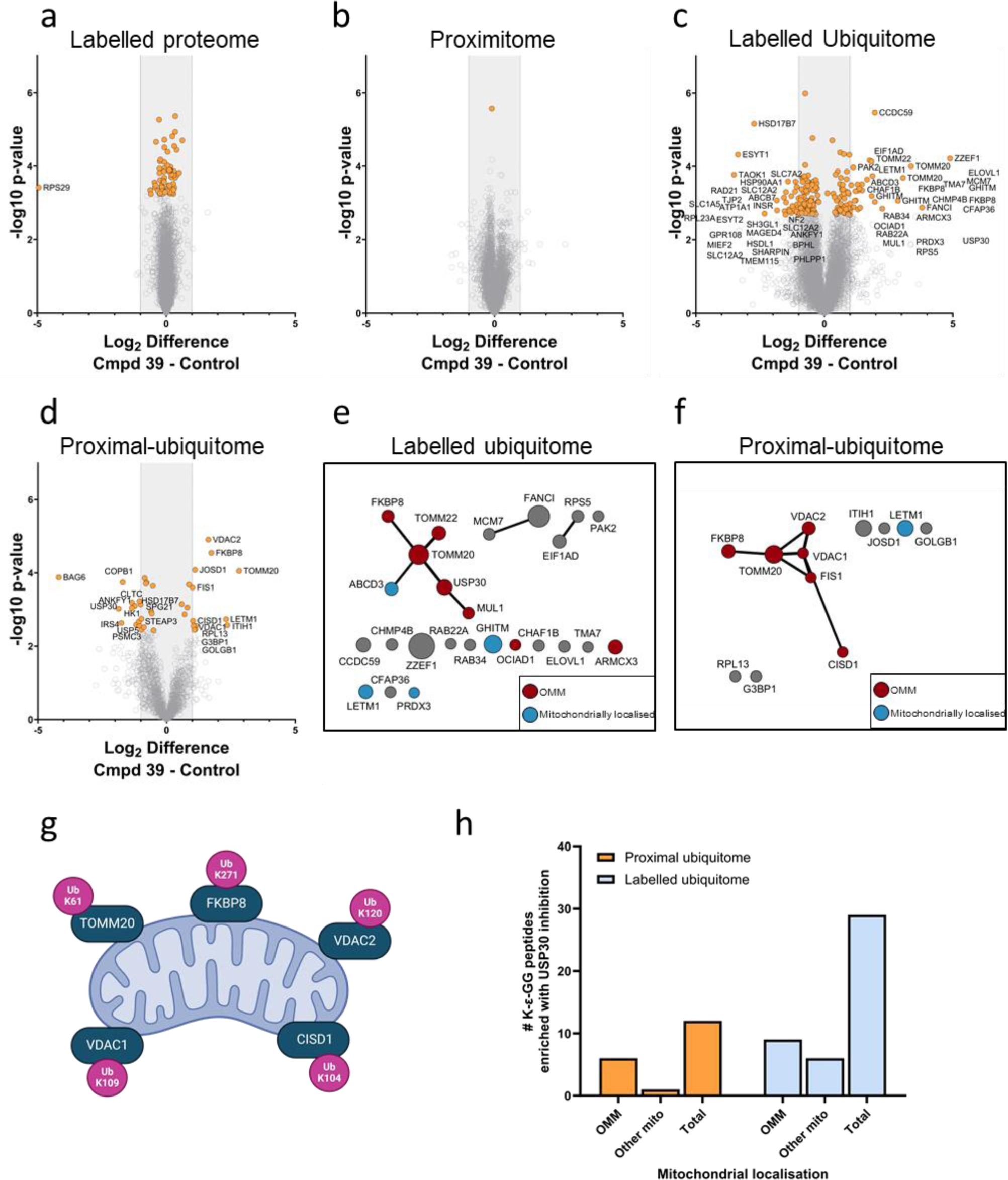
Discovery of USP30 substrate candidates by integrated proximal ubiquitomics. **a.** Labelled proteome volcano plot showing the effect of compound **39** treatment on protein levels in CCCP treated HEK293 USP30-APEX2 cells with APEX2 biotin labelling. **b.** Proximitome volcano plot showing the effect of compound **39** treatment on protein levels captured from streptavidin pulldown after USP30-APEX2 biotinylation in CCCP treated HEK293 USP30-APEX2 cells. **c.** Labelled ubiquitome volcano plot showing the effect of compound 39 treatment on peptides with K-ε-GG sites after K-ε-GG peptide enrichment in CCCP treated HEK293 USP30-APEX2 cells with USP30-APEX2 biotin labelling. **d.** Proximal-ubiquitome volcano plot showing the effect of compound **39** treatment on peptides with K-ε-GG sites after streptavidin pulldown of USP30-APEX2 biotinylated proteins followed by K-ε-GG peptide enrichment in CCCP treated HEK293 USP30-APEX2 cells. **e-f.** Cytoscape string analysis of K-ε-GG peptides that are significantly increased upon compound **39** treatment in CCCP treated HEK293 USP30-APEX2 cells in the labelled ubiquitome **(e)** and proximal ubiquitome **(f)**. Circle size is proportional to the Log_2_ difference in K-ε-GG peptide intensity +/− compound **39** treatment, with the largest difference used if multiple peptides originate from the same protein. Red circles denote OMM and blue circles denote other mitochondrially localised Mitocarta 3.0 annotated proteins. Cytoscape String network based off string confidence score cut off ≥ 0.7. Edge width is proportional to string confidence score.^40,44^ **g.** K-ε-GG sites of the 5 proximal ubiquitome hits that overlap with altered ubiquitination states of proteins with USP30 inhibition/KO reported by Jones *et al.*^22^ and Ordureau *et al.*^24^ **h.** Number of K-ε-GG peptides identified in the proximal ubiquitome and labelled ubiquitome either in total, with OMM localisation or with other mitochondrial localisation annotation (‘other mito’) with Mitocarta 3.0 annotation.

Whilst proteins identified in the labelled proteome and USP30 proximitome were not perturbed upon USP30 inhibition, significant alterations to ubiquitinated peptide intensities as a consequence of USP30 inhibition were observed in the labelled ubiquitome and proximal-ubiquitome (Figure 4c-d). Significant increases in the intensity of ubiquitinated peptides with USP30 inhibition represent potential USP30 substrates. Here, significantly increased ubiquitination (> 1 log_2_ difference) is detected at various sites on 32 different proteins across the labelled ubiquitome and proximal ubiquitome (Figure 4e-f). Of the 32 proteins, 5 overlap between this dataset and two previously reported datasets with significantly increased ubiquitination events in the case of USP30 knockout in depolarizing conditions.^22,24^ Those hits include TOMM20, FKBP8, VDAC2, VDAC1 and CISD1, all of which are Mitocarta3.0 annotated as OMM proteins. Of the 5 hits, 2 are detected as significantly increased with USP30 inhibition in the labelled ubiquitome out of 25 in total. All 5 are detected as significantly increased with USP30 inhibition in the proximal-ubiquitome out of 12 proteins in total (Figure 4g), indicative of the ability of the proximal-ubiquitomics technique to detect robust alterations occurring as a consequence of a reduction in USP30 activity across different cell types and experimental conditions.

The total number of increased ubiquitylation events as a consequence of USP30 inhibition is substantially reduced in the proximal-ubiquitome when compared to the labelled ubiquitome. A greater proportion of increased ubiquitylation events are OMM in the proximal-ubiquitome when compared to the labelled ubiquitome, whereas a smaller proportion of ubiquitylation events are occurring on proteins associated with other parts of the mitochondria (Figure 4h). This finding is reflective of the enrichment of OMM proteins in the proximitome relative to the proteome, and demonstrates that the proximal-ubiquitomics methodology is enabling the detection of changes in ubiquitination events occurring in the proximity of USP30.

### LETM1 is a USP30 interactor and substrate

The increase in ubiquitination of OMM proteins with USP30 inhibition in the proximal-ubiquitome has also been previously reported with USP30 KO.^24^ However, the increased ubiquitination of the inner mitochondrial membrane (IMM) protein LETM1 as a consequence of reduced USP30 activity has not previously been reported to our knowledge. The increase in LETM1 ubiquitination with USP30 inhibition was identified in both the ‘labelled ubiquitome’ and ‘proximal-ubiquitome’, and so LETM1 was further investigated as a potential USP30 substrate.

Initially, we sought to validate the proximity of USP30 to LETM1 using antibody-based proximity ligation assays (PLA), allowing for the detection of proteins within a 30-40 nm proximity via fluorescence. In the absence of a selective USP30 antibody suitable for immunofluorescence (IF) / immunoprecipitation (IP), Flag-USP30 was overexpressed, and PLA was performed both with and without CCCP. A strong PLA signal confirmed that endogenous LETM1 and Flag-USP30 are in proximity regardless of the presence of CCCP (Figure 5a). USP30 has previously been found to Co-IP with TOMM20,^42^ and the increased ubiquitination of TOMM20 with USP30 KO in ubiquitomic experiments has previously been validated by western blot.^22,24^ Therefore, TOMM20 was included as a positive control in experiments investigating LETM1 as a substrate of USP30. Here, endogenous TOMM20 was also identified as proximal to exogenous Flag-USP30 by PLA assays, indicating that the exogenous Flag-USP30 was correctly located on the mitochondria, and co-localised to the TOMM complex (Figure 5b).

**Figure 5.**
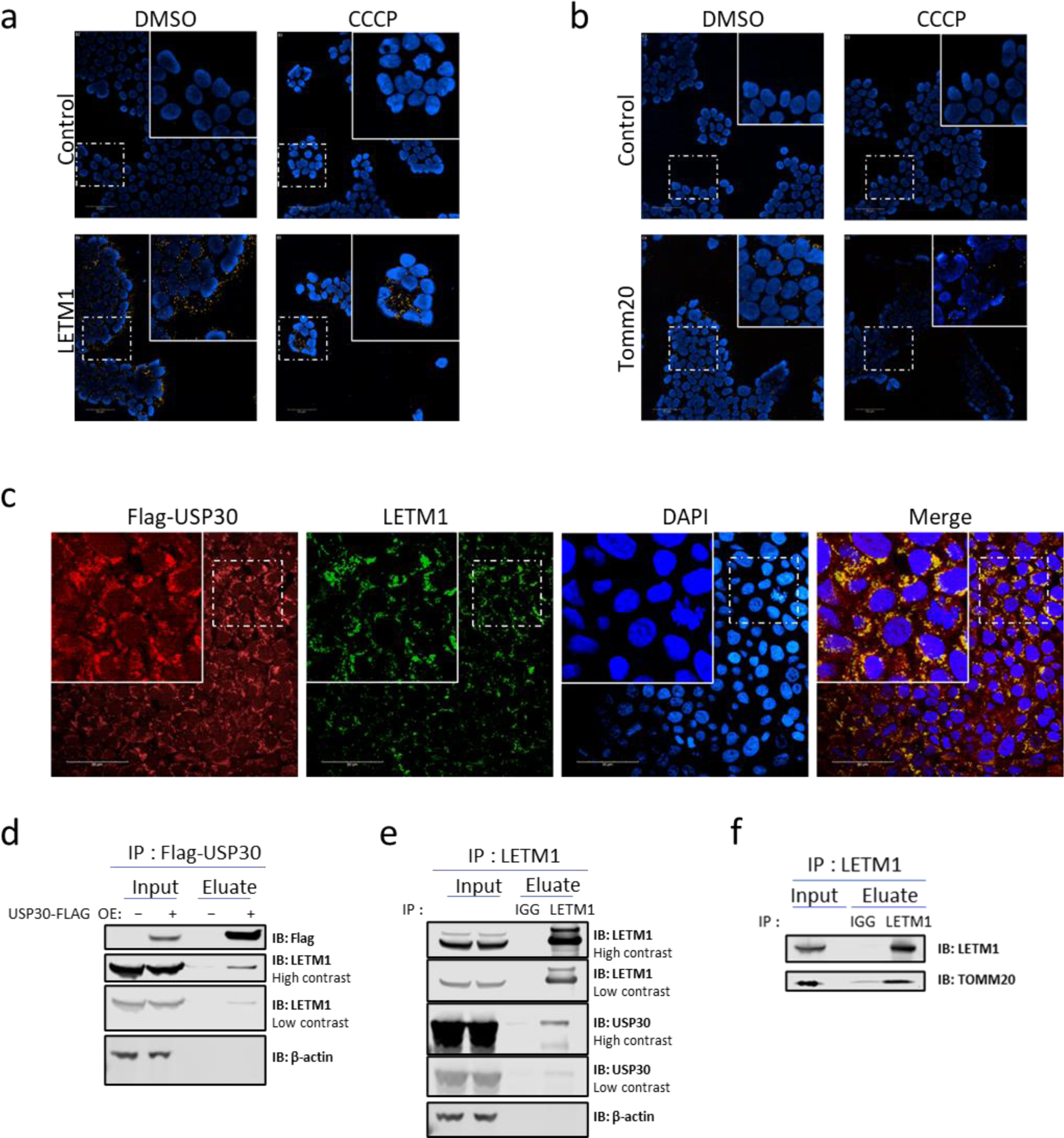
LETM1 as USP30 interactor. **a-b** Proximity Ligation Assay (PLA) in HEK293 cells. HEK293 cells transiently transfected with Flag-USP30, with PLA reaction between anti-Flag antibody and anti-LETM1 antibody **(a)** or an anti-Tomm20 antibody **(b)**. Yellow punctate signals represent the reported interactions, and blue signals indicate nuclei. A single antibody control (None) was included. Scale bar: 10 μm. **c** Confocal images of HEK293 cells +/− CCCP transiently expressing Flag-USP30 and endogenous LETM1 protein. Anti-Flag antibody (red), anti-LETM1 antibody (green), and DAPI (blue) for nuclei visualization. Scale bars: 10 μm. **d-e** HEK293T cells transiently expressing Flag-USP30 **(d)** USP30 was pulled down using an anti-Flag antibody **(e)** LETM1 was pulled down using an anti-LETM1 antibody **(f)** SH-SY5Y cell LETM1 pull down using anti-LETM1 antibody.

To further explore and increase our confidence regarding the proximity of USP30 to LETM1, a co-localization experiment was performed. Flag-USP30 was overexpressed and colocalization with endogenous LETM1 was assessed. Endogenous LETM1 and Flag-USP30 colocalized, supporting the possibility that these two proteins could be in the same protein complex in the presence of CCCP (Figure 5c). To strengthen the validity of our colocalization findings, we calculated Li’s Intensity Correlation Quotient (ICQ) to quantify the degree of correlation between LETM1 and USP30 signals, resulting in a value of 0.182, which is indicative of a positive correlation. The USP30-LETM1 localisation in the presence of CCCP was further validated with Co-IP experiments, where the pulldown of exogenous Flag-USP30 captured endogenous LETM1, and the pull down of endogenous LETM1 captured exogenous Flag-USP30 (Figure 5d-e).

Additionally, Co-IP successfully captured endogenous TOMM20 where endogenous LETM1 was pulled down (Figure 5f). These data collectively support that LETM1 and USP30 are not only proximal to each other but also physically interact, either directly or indirectly. The interaction between these two proteins suggests that LETM1 may be a substrate of USP30, although this does not confirm USP30’s ability to modify LETM1’s ubiquitination status. To further investigate, HEK293 cells were transfected with HA-Ub for 24 hours, treated with CCCP and MG132, and the ubiquitylated proteome was purified by HA pull-down in both the presence and absence of compound **39**. Upon the IP of HA-ubiquitin, LETM1 displayed increased high molecular weight laddering in the presence of compound **39**, suggesting increased LETM1 ubiquitination with USP30 inhibition (Figure 6a). This increased laddering was also phenocopied with USP30 KD (Figure 6b).

**Figure 6.**
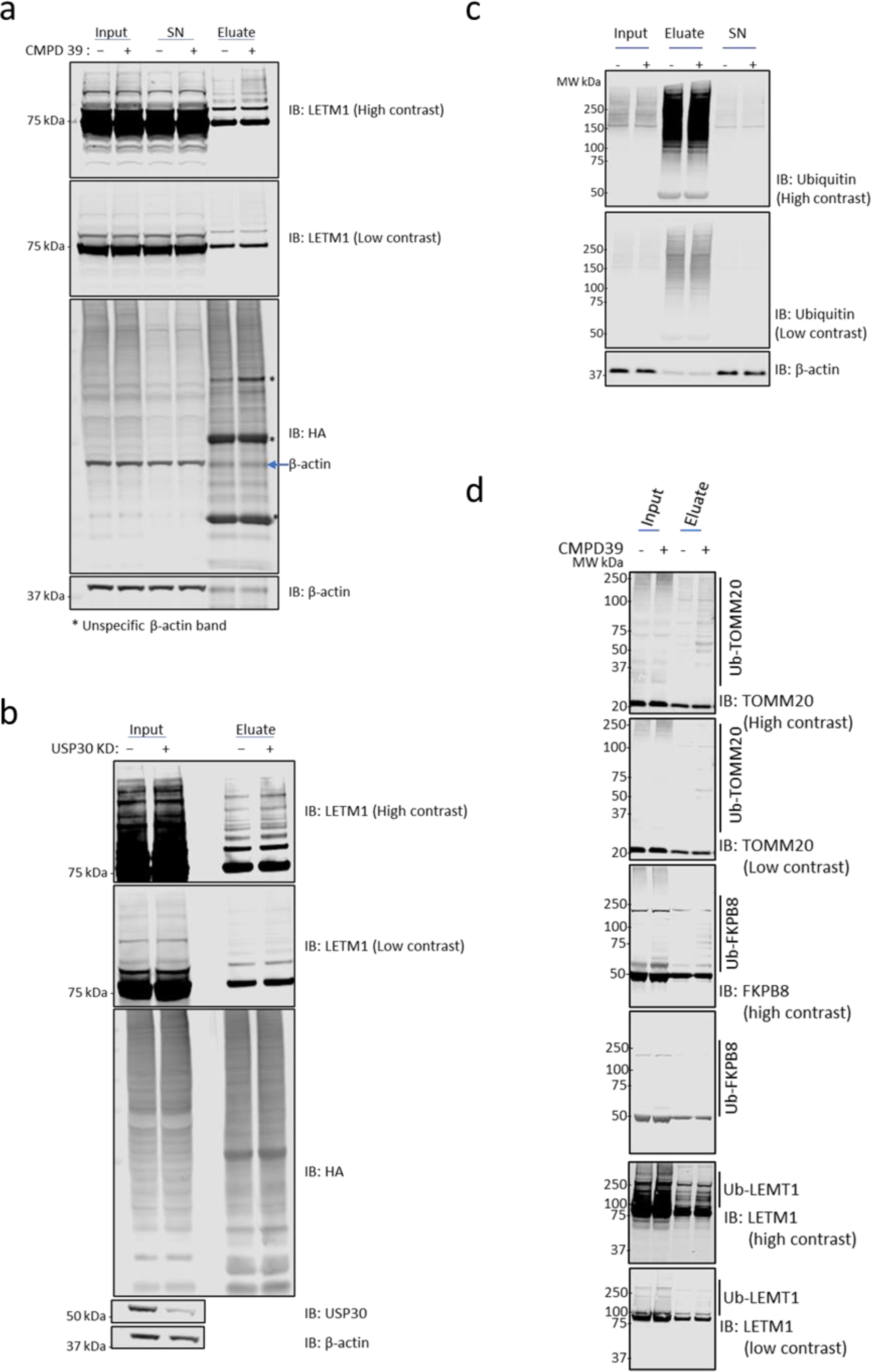
LETM1 and FKBP8 deubiquitination are USP30 dependent. **a** HA-ubiquitin pull down using anti-HA antibody. HEK293T cells transiently expressing HA-Ub treated with 10 µM CCCP +/− Compound 39. **b** HA-ubiquitin pull down using anti-HA antibody. HEK293T cells transiently transfected with scrambled or USP30 siRNA and treated with 10 µM CCCP. (**c-d**) Tandem Ubiquitin Binding Entity (TUBE) pull down in HEK293T cells treated with 10 µM CCCP +/− Compound **39**.

To further validate this ubiquitylation signal with an orthogonal technique, we also performed Tandem Ubiquitin Binding Entity (TUBE) pull downs on HEK293 cells treated with CCCP and +/− compound **39** (Figure 6c). Initially, we assessed the ubiquitination status of TOMM20 as a positive control, observing an accumulation of high molecular weight bands in the USP30-inhibited sample (Figure 6d). An increase in high molecular weight bands with TUBEs enrichment was also seen with FKBP8 (Figure 6d), the ubiquitination of which has previously been reported to increase with USP30 KO,^22,24^ but has not previously been validated.

Finally, upon USP30 inhibition, LETM1 exhibited high molecular weight laddering, indicating its increased ubiquitination in the TUBE-purified ubiquitylated proteome (Figure 6d). Taken together, altered LETM1 ubiquitination is arising as a direct consequence of USP30 activity, and is not due to an enzyme scaffolding function nor an off-target effect of the inhibitor.

## Discussion

The study of optimizing the proximal-ubiquitome of USP30 presented here demonstrates the utility of combining the techniques of proximity labelling and ubiquitomic enrichment to elucidate direct substrates of a DUB/E3 ligase in a cellular environment. Where protein ubiquitination ‘ubiquitomics’ through the enrichment of the K-ε-GG ubiquitin remnant motif can identify global alterations in the ubiquitomic state of proteins, it does not distinguish between direct and downstream ubiquitomic events occurring upon the perturbed activity of a DUB/E3. The dynamic nature of ubiquitination events means that capturing and subsequently validating robust changes directly linked to an enzyme’s activity can be challenging.

Proximity labelling is often employed to profile protein-protein interactions, including transient/weak interactions.^45^ While proximity labelling offers a targeted insight into a protein’s interactions and binding partners, here we demonstrate with APEX2 proximity labelling, that the microenvironment of USP30 is not significantly altered upon inhibition of the enzyme. In this case, it may indicate that the enrichment of those proteins proximal to USP30 alone does not inform on substrates of the enzyme itself. Additionally, high background in proximity labelling experiments means that the proteins identified in the proximitome are not exclusively interactors of a protein of interest.^46^ The power of applying proximity labelling alone rests where large changes in protein localisation or complexes occur as a consequence of a stimulus.

Both ubiquitomic and proximity labelling studies have limitations for the identification of DUB/E3 ligase substrates when applied alone, but, when combined together, they allow for the capture of ubiquitination events proximal to the enzyme of interest. By using ubiquitination as a readout, which changes when comparing an inhibited DUB/E3 ligase to a non-inhibited version, or when comparing a wild-type protein to a catalytic-dead mutant, we overcome limitations of previous methods used to identify substrates of E3 ligases, some of which cannot distinguish substrates from general protein interactors, and none of which can identify specific ubiquitination sites. Additionally, by using APEX2 to label both strong and weakly interacting substrates in the local environment around our bait protein, we overcome the limitation of IP-based methods, which typically only capture strong interactions.

Here, with USP30 as a model system, we have demonstrated that proximal-ubiquitomics enriches for events occurring in the proximity of the enzyme leading to a targeted list of positive ubiquitination hits for further validations. This was demonstrated by follow up experiments exploring LETM1 as a potentially novel USP30 substrate. LETM1 is an IMM protein, which is essential^47^ and is key to multiple mitochondrial functions such as the maintenance of mitochondrial morphology, mitochondrial tubular network assembly and proton-dependent calcium efflux from mitochondria.^30,31,48–51^

Here, upon USP30 inhibition, LETM1 presented with increased ubiquitination at K292, which is proposed to be located in the mitochondrial intermembrane space.^47,51^ We were able to detect the co-localisation of LETM1 to USP30 along with an altered ubiquitination state of LETM1 upon USP30 inhibition and knockdown. USP30 has previously been found to Co-IP with the TOMM complex,^42^ and here we find that LETM1 also Co-IPs with TOMM20. This interaction may be upon mitochondrial import of LETM1 by the TOMM complex, or through interaction within the inter mitochondrial membrane space. Deubiquitination of proteins by USP30 may allow for them to be imported by the TOMM complex.^42^ However, this may not be exactly clear as USP30 deubiquitination protein substrates are observed here to be predominantly located in the OMM, consistent with previous studies.^22^ Further experimentation is needed to fully characterise these processes in greater molecular detail.

The co-localisation of USP30 to the TOMM complex was reproduced here with TOMM20-USP30 PLA and co-localisation, as well as the previously validated increase in TOMM20 ubiquitination with reduced USP30 activity. We also confirmed increased ubiquitination of FKBP8 upon USP30 inhibition, which has been previously reported but not validated. The TOMM complex is critical for the import of mitochondrial preproteins,^52^ and FKBP8 has been identified as mitophagy receptor for Parkin independent mitophagy.^53^ The presence of LETM1, TOMM20 and FKBP8 in the proximal-ubiquitome of USP30 increases the likelihood that they are direct USP30 substrates. Therefore, the proximal-ubiquitomics methodology developed here allows for an advanced insight into the action of the USP30 upon mitochondrial depolarization, and how it may be influencing the mitophagy pathway. The proximal-ubiquitomics approach builds upon previously described E3 ligase substrate detection methodologies, overcoming their limitations and allowing for the identification of specific ubiquitination events that could be applied as robust biomarkers of USP30 inhibition. More generally, proximal-ubiquitomics may be applicable more widely, not only for DUB/E3 substrate discovery, but also to direct PROTAC, molecular glue and DUBTAC candidate molecule design with translational potential.

## Methods

### Cell Culture

Cells were incubated at 37 °C, 5 % CO_2_. HEK293 cells were cultured in high glucose DMEM, supplemented with 10% FBS, and 2% L-glutamine. SH-SY5Y cells were cultured in a 1:1 mix of Hams F12 nutrient Mix and EMEM, supplemented with 15 % FBS, 1 % Non-Essential amino acids and 2 mM Glutamax.

### Plasmid Generation

The USP30-APEX2 was expressed using a pLenti-CMV-BsR-PGK-USP30-APEX-2 plasmid synthesized by Vigene Biosciences. APEX2 was conjugated to the C terminal of USP30, with a glycine/serine flexible linker. Flag-HA-USP30 was a gift from Wade Harper (Addgene plasmid # 22578; http://n2t.net/addgene:22578; RRID:Addgene_22578).^54^

### Lentiviral packaging

HEK293T cells were plated in a T175 flask and incubated at 37 °C with 5 % CO_2_ until they reached 85% confluency. The cells were then transfected with pMD2G, psPAX2, and the USP30-APEX2 plasmids using Lipofectamine 2000 according to the manufacturer’s protocol. After two days of incubation, the supernatants were collected, filtered through a 0.45 μm filter, and subjected to ultracentrifugation to concentrate the viral particles.

### Cell line Generation

1.5×10⁵ HEK293 cells were seeded in a 6-well plate and incubated overnight at 37 °C and 5 % CO_2_. 24 hrs after seeding, USP30-APEX2 lentivirus with polybrene was added to the HEK293 cells at a concentration of 8 µg/ml and incubated overnight at 37 °C and 5 % CO_2_. 24 hrs after lentiviral treatment, the HEK293 cells were trypsinized and seeded into a 100mm dish (All cells were used). After 8 hours, 5 μg/ml blasticidin was added to both the negative control and the transduced cells. The cells were incubated for 10 days under selection, with the medium being replaced every 3 days.

### Transient Transfection

For DNA plasmid transfections, cells were seeded in 6-well plates and transfected for 24 hrs using Lipofectamine 3000 Transfection Reagent (L3000001, ThermoFisher UK) according to the manufacturer’s instructions. The concentration of plasmids used can be found in the details of each specific experiment.

For siRNA transfection experiments, cells were grown in 6-well plates and transfected for 72 hrs with RNAiMAX Transfection Reagent (Invitrogen, 13778-150) following the manufacturer’s protocol. A final concentration of 10 nM was used for the following Silencer Select siRNAs: control siRNA (4390844) and si-USP30 (Dharmacon ON-TARGETplus SMARTpool).

### Compound 39 synthesis

‘compound **39**’, refers to CAS number 2242582-40-5. The inhibitor was >95% pure as assessed by HPLC analysis.^27^

### HA-Ub-PA ABP labelling

Ubiquitin (Gly76del) with an N-term HA tag and C-term propargylamine (HA-Ub-PA) synthesis and lysate labelling was carried out as previously described.^26^ HEK293 expressing USP30-APEX2 were treated with compound **39** for 6 hrs at the concentration indicated in the figure, washed 3 times in PBS and lysed in 50 mM Tris (pH 7.5), 5 mM MgCl_2_, 0.5 mM EDTA, 250 mM sucrose and 1 mM DTT with bead beating. Lysates were then clarified at 600 x*g* for 10 minutes at 4 °C and supernatant protein was quantified by BCA. HA-Ub-PA was then incubated with lysate protein (1:100 w/w) for 5 minutes at 37 °C.

### Membrane-Cytoplasmic Cellular Fractionation

As previously described^39^, fresh HEK293 FRT cell pellets were resuspended in digitonin lysis buffer (50 mM Hepes, pH 7.5, 10 mg/ml digitonin, 150 mM NaCl) and incubated for 30 minutes at 4°C with continuous agitation. The sample was then centrifuged at 6,000g for 5 minutes at 4°C. The resulting supernatant, containing the cytoplasmic fraction, was carefully transferred to a new tube. The pellet was resuspended in a buffer containing 0.3% NP-40, 50 mM Hepes (pH 7.5), and 150 mM NaCl, and incubated on ice for 5 minutes. After centrifugation at 1,500g for 5 minutes, the supernatant, representing the membrane fraction, was collected into a separate tube and stored at −20°C until further analysis.

### Proximal ubiquitomics workflow - APEX2 proximity labelling

HEK293 cells expressing USP30-APEX2 were grown in T175 flasks coated with Poly-D-Lysine to 100 % confluency, with 5 x T175 flasks pooled per condition. The medium was then replaced with fresh medium containing 10 µM CCCP, with either 5 µM of compound **39**, or DMSO as a control. The cells were incubated at 37°C and 5% CO_2_ for 5.5 hours. Following this incubation, proximity labelling was performed as previously described.^34,46,55^ Briefly, 2 µM phenol biotin was added to the media, and the cells were incubated for an additional 30 minutes. Subsequently, H_2_O_2_ at a concentration of 1 mM was added to the medium for exactly 1 minute. The cells were then washed four times with Quencher buffer (10 mM sodium ascorbate, 10 mM sodium azide, 5mM Trolox in DPBS), with each wash involving a 1-minute incubation.

The cells were lysed using lysis buffer (10 ml per 5 x T175 flasks) (RIPA lysis buffer containing 1 mM PMSF, 5 mM Trolox, 10 mM sodium azide, 10 mM sodium ascorbate, PhosSTOP phosphatase inhibitor and cOmplete Protease Inhibitor Cocktail) and centrifuged at 10,000 x*g* for 10 minutes at 4°C. At this stage 5% of lysate was taken for the ‘labelled proteome’ and processed for mass spectrometry analysis using the S-Trap™ micro digestion protocol according to the manufacturer’s instructions. and the remain lysate was incubated with 3.5 mL of streptavidin agarose beads for 1 hour at room temperature. The beads were then washed with 10 mL of the following buffers in the specified order: twice with RIPA buffer, then with KCl, Na_2_CO_3_, 2M Urea, 1% SDS and finally with 2 times with RIPA buffer.

Biotinylated proteins were eluted in 4 mL of 2x loading dye supplemented with 2 mM biotin and 20 mM DTT by boiling (98°C) for 10 min. Eluted material was subsequently processed for mass spectrometry analysis using the S-Trap™ midi digestion protocol according to the manufacturer’s instructions. 10 % of eluates from the S-Trap™ clean up were taken for LC-MS/MS analysis as the ‘proximitome’.

### Proximal ubiquitomics workflow - K-ε-GG immunoprecipitation

The subsequent K-ε-GG immunoprecipitation (IP) was performed according to the manufacturer’s instructions with small differences. Briefly, lyophilized peptides were centrifuged for 5 minutes at 2000 x*g* at room temperature. To the dried peptides, 0.5 mL of 1X IAP bind buffer was added, and the peptides were resuspended mechanically by repeated pipetting. The solution was then centrifuged for 5 minutes at 10,000 x*g* at 4°C and transferred into a new Eppendorf tube.

For each sample, 10 µL of bead slurry was placed into a 1.5 mL Eppendorf tube. The beads were washed four times with 1 mL of ice-cold PBS. The soluble peptides were then transferred to the beads and incubated on an end-over-end rotator for 2 hours at 4°C. Afterwards, the samples were spun down at 2000 x*g* for 5 seconds to pellet the beads, placed on a magnetic rack for 10 seconds, and the unbound peptide solution was discarded.

The beads were washed with 1 mL of chilled HS IAP Wash Buffer (1X), resuspended, briefly centrifuged as before, placed on the magnetic rack for 10 seconds, and the wash buffer was removed. This washing step was repeated three additional times. The samples were then washed twice with 1 mL of chilled LCMS water. Ubiquitinated peptides were eluted by adding 50 µL of IAP Elution buffer and incubating at room temperature for 10 minutes with gentle mixing. The elution was repeated once more to ensure complete recovery of ubiquitinated peptides. Peptides were then purified by C18 and resuspended in 5% DMSO/5% FA for LC-MS/MS analysis.

The input material optimization shown in Figure 2g-h follow this USP30-APEX2 proximity labelling workflow with indicated number of 15 cm dishes as input.

### Control proteome, control ubiquitome and labelled ubiquitome

HEK293 cells expressing USP30-APEX2 were grown in 10 cm dishes coated with Poly-D-Lysine to 100 % confluency, with 1 x 10 cm dish used per condition. Cells were APEX2 labelled as detailed for the proximal-ubiquitome for the ‘labelled ubiquitome’. Cells for the control proteome and control ubiquitome were not APEX2 labelled. Cells were treated with CCCP +/− compound **39**, lysed in 500 µL, clarified and processed using S-Trap™ midi as detailed for the proximal-ubiquitome. Where cells were not APEX2 labelled, 5% was removed for the control proteome. Remaining peptides were then K-ε-GG immunoprecipitated as detailed for the proximal-ubiquitome.

### Liquid chromatography tandem mass spectrometry - LC-MS/MS

The proximal-ubiquitome was analysed on a timsTOF SCP (Bruker) LC-MS/MS system to maximise sensitivity. The input material optimization detailed in Figure 2g-h was analysed on an Orbitrap Fusion Lumos Tribid (Thermo) LC-MS/MS system. All other samples were analysed on an Orbitrap Ascend Tribid (Thermo) LC-MS/MS system.

APEX2 proximitome and total proteomics samples were analysed by LC-MS/MS using a Vanquish Neo UHPLC (Thermo) connected to a Thermo Orbitrap Ascend mass spectrometer (Thermo). The Vanquish Neo was operated in “Trap and Elute” mode using a PepMap Neo trap (185um, 300um x 5mm) and EASY-SPRAY PepMapNeo column (50cmx75um, 1500bar). Tryptic peptides were trap and separated using a 60 min linear gradient over 60 minutes (from 3 to 20% B in 40 min and from 20 to 35% in 20 min) at 300nl/min flow.

APEX proximitome and total proteomics samples were acquired in DIA mode with some slight changes from previously described method.^27,56^ Briefly, survey scans (MS1) were acquired in the Orbitrap over the mass range of 350 – 1650m/z, with a 45k resolution, maximum injection time of 91 ms, an AGC set to 125% and a RF lens at 30%. MS2 scans were then collected using the tMSn scan function, with a 40 predefined variable width DIA scan windows^56^ with a 30K orbitrap resolution, normalized AGC target of 1000%, maximum injection time set to auto and a 30 % collision energy.

K-ε-GG enriched ubiquitomics samples were analysed by LC-MS/MS using an Ultimate 3000 HPLC coupled to an Orbitrap Ascend Tribrid or an Orbitrap Fusion Lumos Tribid instrument (Thermo Fisher) using a nano-EASY spray source. Tryptic peptides were loaded onto an AcclaimPepMap100 trap column (100µm x 2cm, PN164750; ThermoFisher) and separated on a 50cm EasySpray column (ES903, ThermoFisher) using a 60 min linear gradient from 2 to 35% acetonitrile, 0.1% formic acid and at 250nl/min flow rate. Both trap and column were kept at 50C. Data were acquired in DIA-mode with 44 variable width windows optimised for GG workflows as previously described.^37^ In brief, MS1 scans were acquired in the Orbitrap over the mass range of 350 – 1650m/z, with a 120k resolution, maximum injection time of 60ms, an AGC set to 300% and a RF lens at 30%. MS2 scans were then collected using the tMSn scan function, with a 44 predefined variable width DIA scan windows^37^ with a 30K orbitrap resolution, normalized AGC target of 2000%, maximum injection time set to auto and HCD stepped collision energy set at 22, 27 and 30 %.

Proximal-ubiquitome samples were analysed using an Evosep One LC system (EvoSep) coupled to a timsTOF SCP mass spectrometer (Bruker) using the Whisper 40 samples per day method and a 75 µm x 150 mm C18 column with 1.7 µm particles and an integrated Captive Spray Emitter (IonOpticks). Buffer A was 0.1% formic acid in water, Buffer B was 0.1% formic acid in acetonitrile. Data was collected using diaPASEF^57^ with 1 MS frame and 9 diaPASEF frames per cycle with an accumulation and ramp time of 100 ms, for a total cycle time of 1.07 seconds. The diaPASEF frames were separated into 3 ion mobility windows, in total covering the 400 – 1000 m/z mass range with 25 m/z-wide windows between an ion mobility range of 0.64–1.4 Vs/cm^2^. The collision energy was ramped linearly over the ion mobility range, with 20 eV applied at 0.6 Vs/cm^2^ to 59 eV at 1.6 Vs/cm^2^.

### LC-MS/MS data processing

All data were searched library-free with DIA-NN 1.8.1^37,58^ with a Uniprot *Homo Sapiens* fasta (20,370 entries, retrieved on April 16, 2021). For ubiquitome searches all setting were left as default other than the addition of variable modifications (methionine oxidation, N-terminal acetylation and K-ε-GG ubiquitin remnant motif, with a maximum of 2 variable modifications allowed). Default search settings include 1 missed Trypsin/P cleavage, with fixed N-terminal methionine excision and fixed cysteine carbamidomethylation. For proteome and proximitome searches, settings were identical other than the K-ε-GG variable modification. Match-between runs was enabled for all searches, except for the input material optimization in Figure 2g-h.

### LC-MS/MS data analysis

For ubiquitome data, peptide level information was extracted from the DIA-NN precursor matrix output. The output was filtered to remove any non-proteotypic identifications, collapsed to provide peptide level information by summing precursors with different charge states using R (version 4.3.2), and filtered for the UniMod:121 K-ε-GG adduct. For proximitome and proteome data, the DIA-NN protein groups output matrix was used. All data was imported into Perseus (Version 1.6.15.0),^59^ log_2_ transformed, filtered so that values were present in all replicates for at least 1 condition, median subtract normalised, and imputed using imputeLCMD k-nearest neighbour (KNN) with a neighbour number of 15. A Perseus two-sample students T-test with permutation-based FDR set to 5 % was applied to extract significant changes as indicated in figures.

### Permutation analysis of proximitome vs proteome

Permutation analysis was carried out in R to establish the significance of the increased intensity of OMM proteins in the proximitome vs the proteome, when compared to all proteins and to those in the MM. This was achieved by multiplying the difference of each protein between the proximitome *vs* the proteome randomly by either +1 or −1 100000 times and then plotting the means of the differences as a distribution and comparing this to the mean difference of Mitocarta 3.0 annotated OMM proteins, MM proteins and to the proteins overall.

### Immunofluorescence/Proximity ligation assay

Cells were fixed with 4% paraformaldehyde at room temperature for 10 minutes, followed by three washes with PBS. Next, the cells were incubated with PBS containing 0.1% Triton-X-100 for 10 minutes at room temperature, followed by a PBS wash for 5 minutes. Subsequently, the cells were blocked with 5% BSA in TBST containing 0.1% Tween 20 for 1 hour.

Depending on the sample, the cells were then incubated with the following primary antibodies: ANTI-FLAG® M1 (1:1000) (Sigma Aldrich F3040), LETM1 (1:100) (Proteintech, 16024-1-AP), and TOMM20 (1:100) (Cell Signaling, 42406) at 4 °C overnight. After primary antibody incubation, the cells were washed three times with TBST containing 0.1% Tween 20, incubating with the wash buffer for 5 minutes each time.

For the proximity ligation assay (PLA), the cells were incubated with PLA anti-mouse PLUS and anti-rabbit MINUS Duolink secondaries, followed by Duolink In Situ Orange Reagents, according to the manufacturer’s protocol (Millipore Sigma).

For immunofluorescence, the cells were incubated with the secondary antibodies (Goat anti-Mouse IgG (H + L), Alexa Fluor 488-conjugated, and Goat anti-Rabbit IgG (H + L), Alexa Fluor 633-conjugated) in 1% BSA in TBST containing 0.1% Tween 20 for 1 hour at room temperature. The cells were then washed three times with TBST containing 0.1% Tween 20, incubating with the wash buffer for 5 minutes each time.

For nuclear staining, the cells were incubated with Hoechst 33342 (1:1000) for 10 minutes at room temperature, followed by three washes with PBS.

Samples from both PLA and immunofluorescence were imaged using a high-content laser-based spinning disk confocal microscope (Opera Phenix Plus) with a 40x water objective. Images were collected and analyzed using Harmony® imaging and analysis software.

For colocalisation analysis, the images were processed using ImageJ, and Li’s Intensity Correlation Quotient (ICQ) was calculated using the Coloc 2 plugin.

### Immunoprecipitation

#### IP-Flag

Cells were washed twice with ice-cold PBS and then lysed in Pierce IP lysis buffer (25 mM Tris-HCl pH 7.4, 150 mM NaCl, 1% NP-40, 1 mM EDTA, 5% glycerol, PhosSTOP phosphatase inhibitor, and cOmplete Protease Inhibitor Cocktail). The lysates were incubated on ice for 10 minutes with periodic mixing and then clarified by centrifugation at 13,000 x*g* for 10 minutes at 4°C. The pre-cleared lysates were incubated with anti-FLAG M2 Magnetic Beads at 4°C for 4 hours. After washing three times with lysis buffer, the proteins bound to the anti-FLAG antibody were pulled down by 3X FLAG peptide elution buffer. Both the whole cell lysates and the pull-down samples were subjected to immunoblotting analysis.

#### IP-LETM1

Cells were washed twice with ice-cold PBS and then lysed in Pierce IP lysis buffer (25 mM Tris-HCl pH 7.4, 150 mM NaCl, 1% NP-40, 1 mM EDTA, 5% glycerol, PhosSTOP phosphatase inhibitor, and cOmplete Protease Inhibitor Cocktail). The lysates were incubated on ice for 10 minutes with periodic mixing and then clarified by centrifugation at 13,000 g for 10 minutes at 4°C to remove cell debris. The pre-cleared lysates were incubated with anti-LETM1 antibody in a dilution of 1:100. The next day, lysates were incubated with 50uL of Dynabeads™ Protein G for Immunoprecipitation and incubated for 30 minutes at room temperature. After washing three times with lysis buffer, the proteins bound to the anti-LETM1 antibody were eluted by 2X Loading dye and 100mM DTT elution buffer. Both the whole cell lysates and the pull-down samples were subjected to immunoblotting analysis.

### Tandem ubiquitin binding entity pulldown

HEK293 cells were grown in 150 mm dish to 100% confluency, with 1 x 150 mm dish per condition. The medium was then replaced with fresh medium containing 10 µM CCCP, with either 5 µM of compound **39**, or DMSO as a control. The cells were incubated at 37 °C and 5 % CO_2_ for 6 hours. Cells were washed twice with ice-cold PBS and then lysed in Pierce IP lysis buffer (25 mM Tris-HCl pH 7.4, 150 mM NaCl, 1% NP-40, 1 mM EDTA, 5% glycerol, PhosSTOP phosphatase inhibitor, and cOmplete Protease Inhibitor Cocktail, 100mM N-Ethylmaleimide (NEM) and 10 µM MG132). The lysates were incubated on ice for 10 minutes with periodic mixing and then clarified by centrifugation at 13,000 g for 10 minutes at 4°C to remove cell debris. Supernatant was collected and 10 % of sample was removed as “INPUT” for analysis by Western blotting. Endogenous ubiquitylated proteins were isolated from the soluble lysate at 4 °C for 12 hrs using 50 µl of packed agarose TUBE (TUBE1, Life sensors, performed according to the manufacturer’s instructions). Following four washes in TBST, bound proteins were eluted using reducing and denaturing SDS sample loading buffer and analysed by Western blot.

## Data availability

The mass spectrometry proteomics data have been deposited to the ProteomeXchange Consortium via the PRIDE^60^ partner repository with the dataset identifier PXD054890.

## Acknowledgements

We thank members of the Kessler and ODDI groups for helpful discussions. We also thank Val Miller and Daniel Ebner for using and guiding us with confocal microscopy. We would like to express our gratitude to Jesper Hansen for his guidance in analyzing the colocalization experiments. We thank Daryl S. Walter from Evotec for providing us with an aliquot of compound 39. H.B.L.J. was supported by a Bristol-Myers Squibb fellowship. A.D., S.D., and B.M.K. were supported by the Chinese Academy of Medical Sciences (CAMS) Innovation Fund for Medical Science (CIFMS), China [grant number: 2018-I2M-2-002], and B.M.K by the Engineering and Physical Sciences Research Council (EPSRC) grant [number: EP/N034295/1].

## Author contributions

A.D and H.B.L.J contributed equally. A.D., H.B.L.J and B.M.K conceptualized the study. A.D, H.B.L.J, A.G., I.V. and S.D. conducted experiments. H.B.L.J and A.D. analysed data. B.M.K. supervised the experimental work. Funding was acquired by B.M.K. All authors wrote and approved the paper.

