## Supplementary Figures for "Integrative Proximal-Ubiquitomics Profiling for Deubiquitinase and E3 Ligase Substrate Discovery Applied to USP30"

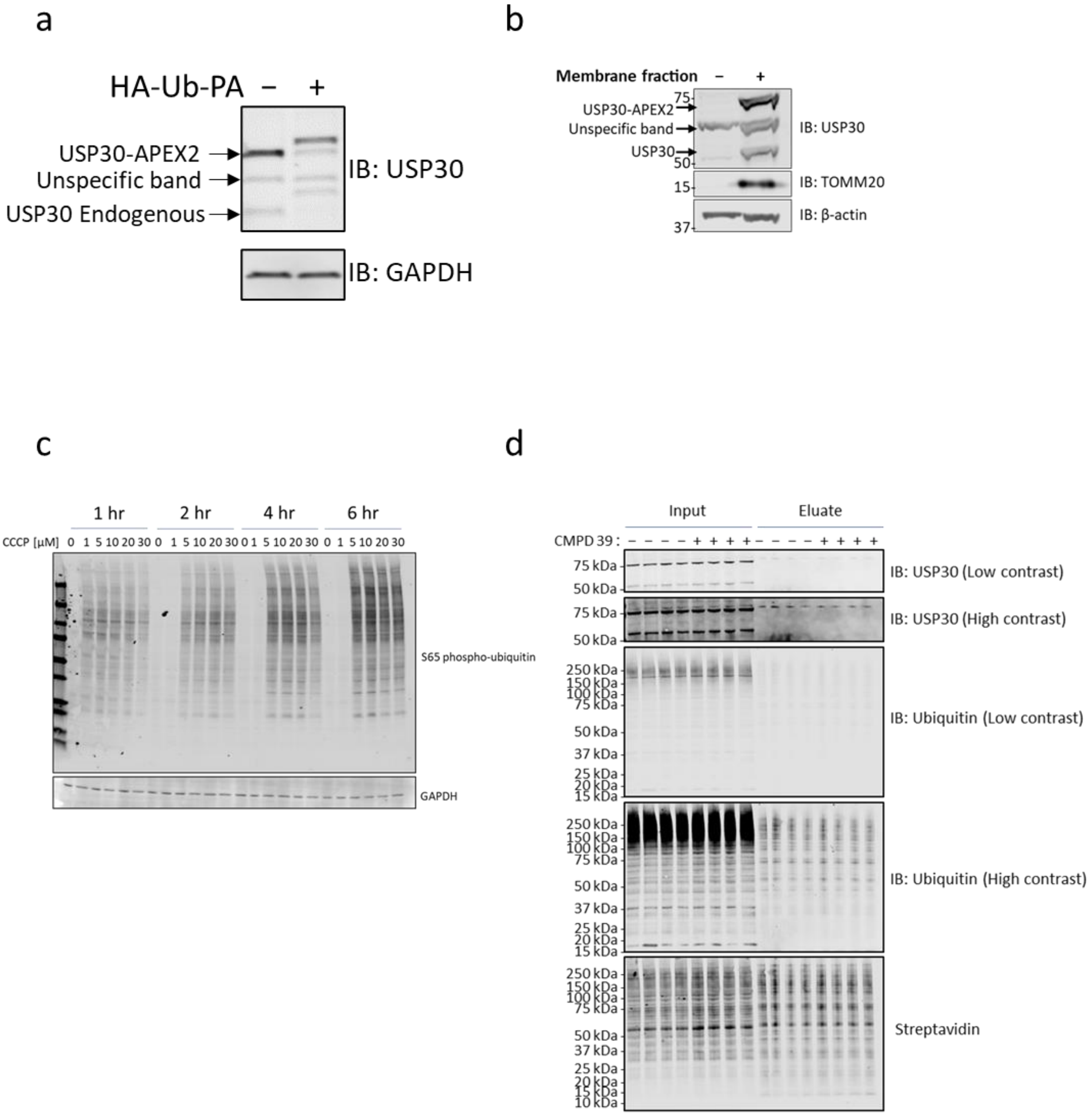


**Figure S1 | a.** USP30-APEX2 expression vs endogenous USP30 expression, +/− HA-Ub-PA labelling. **b.** USP30-APEX HEK293 digitonin membrane separation. **c.** USP30-APEX2 HEK293 S65 phospho-ubiquitin signal with CCCP time and concentration dependence. **d.** Input and elution of all replicates of streptavidin biotin-phenol pulldown.

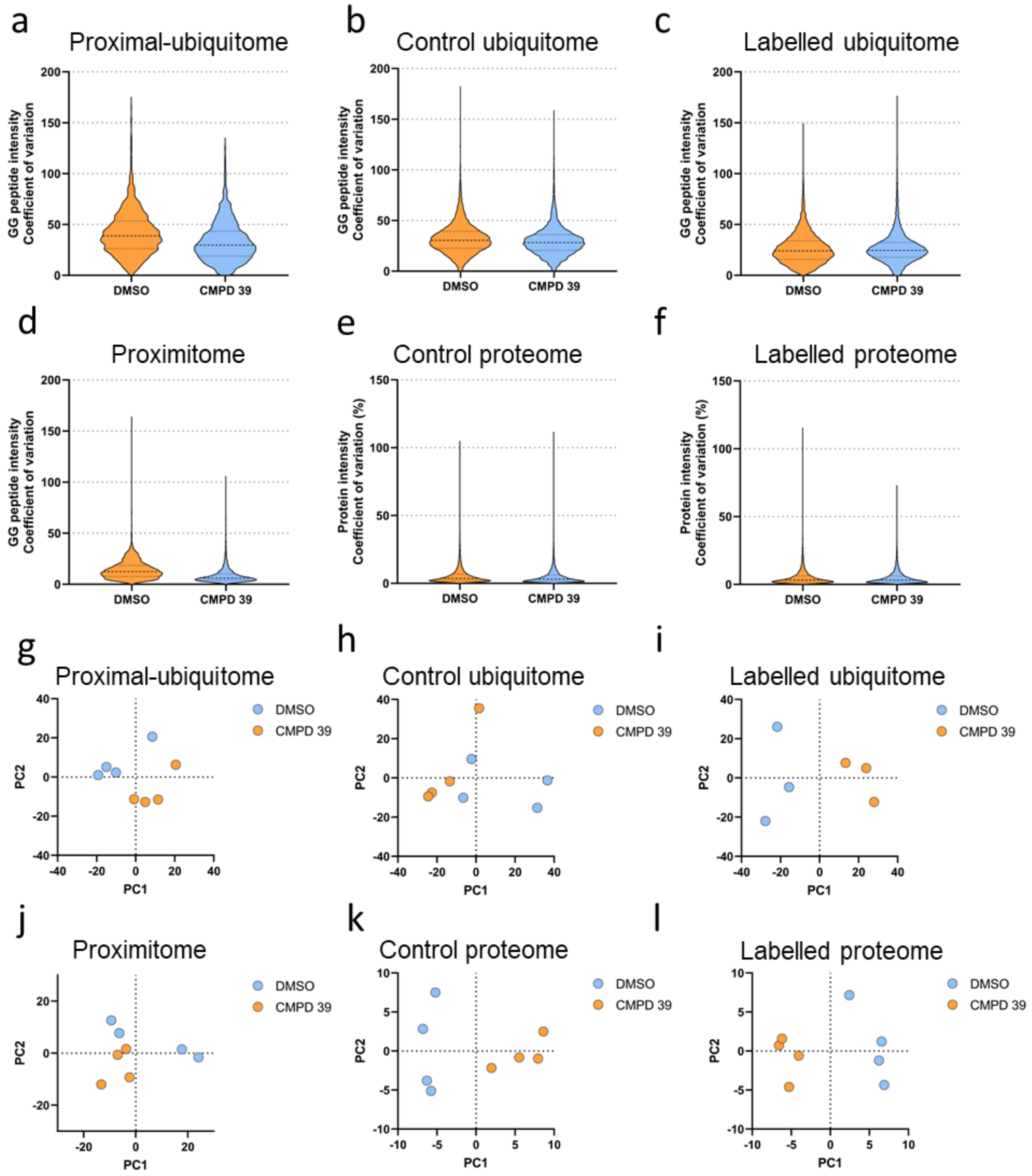


**Figure S2 | a-d** Coefficient of Variation for the proximal ubiquitome (a), the control ubiquitome (b), the labelled ubiquitome (c), the proximitome (d), the control proteome (e), and the labelled proteome (f). **g-l** Principal component analysis of all samples for the proximal ubiquitome (g), the control ubiquitome (h), the labelled ubiquitome (i), the proximitome (j), the control proteome (k), and the labelled proteome (l).

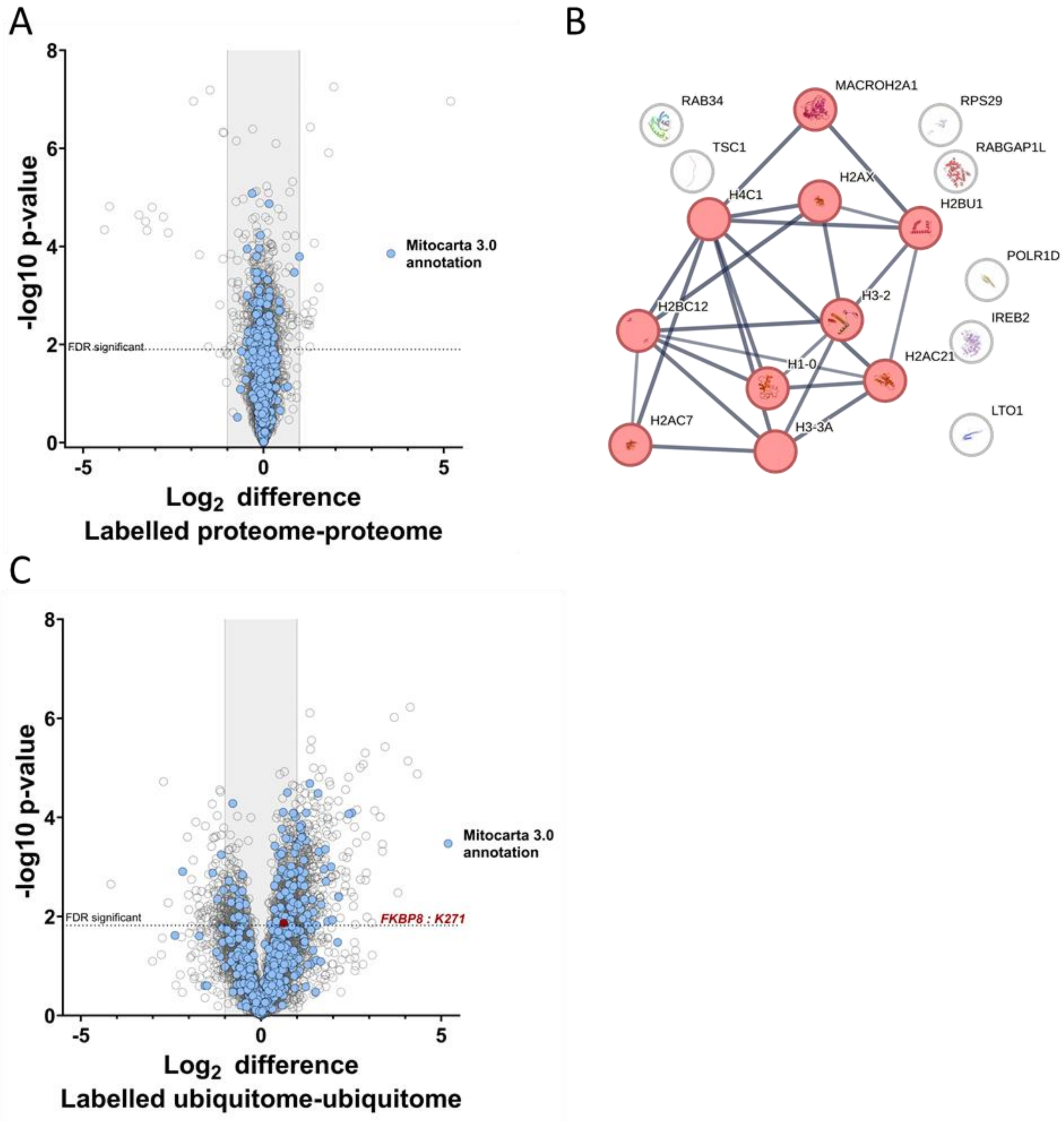


**Figure S3 | a.** Volcano plot of protein intensities from the DMSO treated labelled proteome vs the control proteome. **b.** String analysis of proteins significantly increased in the labelled proteome compared to the control proteome. Proteins in red represent a cluster of 'Structural constituents of chromatin'. Confidence score cutoff ≥ 0.7. Edge width is proportional to string confidence score.[33,37] **c.** Volcano plot of K-ε-GG peptides intensities from the DMSO treated labelled ubiquitome *vs* the control ubiquitome. Significantly altered intensities that overlap with increased ubiquitination events as a consequence of USP30 inhibition in the proximal ubiquitome are highlighted in red.
